# Methionine availability influences essential H3K36me3 dynamics during cell differentiation

**DOI:** 10.1101/2023.11.22.568331

**Authors:** Yudong Sun, Vijyendra Ramesh, Fangchao Wei, Jason W. Locasale

## Abstract

Histone modifications are integral to epigenetics through their influence on gene expression and cellular status. While it’s established that metabolism, including methionine metabolism, can impact histone methylation, the direct influence of methionine availability on crucial histone marks that determine the epigenomic process remains poorly understood. In this study, we demonstrate that methionine, through its metabolic product, S-adenosylmethionine (SAM), dynamically regulates H3K36me3, a cancer-associated histone modification known to influence cellular status, and myogenic differentiation of mouse myoblast cells. We further demonstrate that the methionine-dependent effects on differentiation are mediated in part through the histone methyltransferase SETD2. Methionine restriction leads to preferential decreases in H3K36me3 abundance and genome accessibility of genes involved in myogenic differentiation. Importantly, the effects of methionine restriction on differentiation and chromatin accessibility can be phenocopied by the deletion of Setd2. Collectively, this study demonstrates that methionine metabolism through its ability to be sensed by chromatin modifying enzymes can have a direct role in influencing cell fate determination.

## Introduction

Histone modifications, such as the methylation of lysine residues, constitute a fundamental epigenetic mechanism, fine-turning gene expression and offering an additional layer of control in cell fate determination^1–4^. A pronounced interplay exists between histone modifications and cellular metabolism^5–7^. Enzymatic reactions responsible for the placement and removal of histone modifications, such as acetylation and methylation, rely on substrates derived from core metabolic pathways. For instance, acetylation utilizes acetyl-CoA from the glycolysis / TCA cycle^8^, while methylation involves S-adenosylmethionine (SAM) from the methionine cycle^9^. Further underscoring this nexus, metabolic intermediates from the TCA cycle, such as α-ketoglutarate (α-KG) produced and consumed by the isocitrate dehydrogenase (IDH), serve as cofactors for histone demethylases that catalyze the removal of methylation marks^10^. Moreover, 2-hydroxyglutarate (2-HG), a product of mutated IDH, acts as an inhibitor of histone demethylases^11,12^. These connections illustrate a link between metabolic pathways and epigenetic regulators and open the possibility of modulating biologic processes associated with changes to the epigenetic landscape with metabolic interventions, such as diet.

While several studies have delved into the complexities of how 2-HG/α-KG and histone demethylases influence histone methylation marks and cellular processes, notably differentiation^13–15^, the reciprocal but equally essential process of methylation mediated by histone methyltransferases remain comparatively underexplored. Previous work from our group and others has revealed that methionine and SAM availability can affect critical histone methylation marks like H3K4me3^9,16^. Yet, how methionine restriction affects cellular differentiation, and the dynamics between methionine availability and specific histone methylation marks is unknown.

In this study, we explored the effect of methionine restriction on H3K36me3 dynamics and its subsequent influences on cellular differentiation in cultured mouse myoblast cells. Our findings revealed a dependency of H3K36me3 levels and myogenetic differentiation on methionine, with a reversible character indicative of dynamic regulation. Moreover, our data show that the impact of methionine on differentiation operates, at least in part, through the histone methyltransferase SETD2, establishing it as a sensor for methionine in this context. Additionally, methionine restriction appears to preferentially reduce H3K36me3 abundance and genome accessibility at genes related to muscle differentiation, further bolstering support of a metabolic-epigenetic crosstalk in the regulation of chromatin dynamics and cellular differentiation through methionine and histone methylation.

## Results

### Myogenic differentiation and H3K36me3 are methionine-dependent

C2C12 cells initiate myogenic differentiation when transitioning from growth media (GM) to differentiation media (DM) upon reaching confluence^17^. To investigate the impact of methionine availability on this differentiation process, cells were cultured to confluence in GM containing full methionine (MET) at 200 µM. Upon transition to DM, we either restricted methionine (MR) to 20 µM or depleted it entirely (MD, 0 µM) (Figure 1A). Throughout a five-day differentiation window, cells subjected to MR or MD displayed disruption in differentiation, characterized by a lack of elongated myotube formation (Figure 1B). Further, we developed a machine learning-based tool to quantify differentiation from phase-contrast microscope images (Methods) and observed reduced differentiation for cells on MR and MD conditions at D3 and D5 relative to controls (Figure 1C). Impaired differentiation under MR and MD conditions was further validated by assessing protein levels of myosin heavy chain (MyHC) by immunoblotting (Figure 1D). Concurrent assessment of methionine cycle intermediates (Figure 1E) confirmed the marked reduction in their abundance under MR or MD (Figures 1F - I). In exploring the epigenetic ramifications of these findings, we assessed several key histone methylation marks. Of these, H3K36me3 showed the most pronounced reduction in response to both MR and MD (Figure 1J). Chromatin Immunoprecipitation Sequencing (ChIP-Seq) analysis for H3K36me3 indicated that while its abundance across all genes remained relatively stable during the differentiation period (Figure 1K, Methods), MR markedly reduced its levels (Figures 1L, 1M).

**Figure 1:**
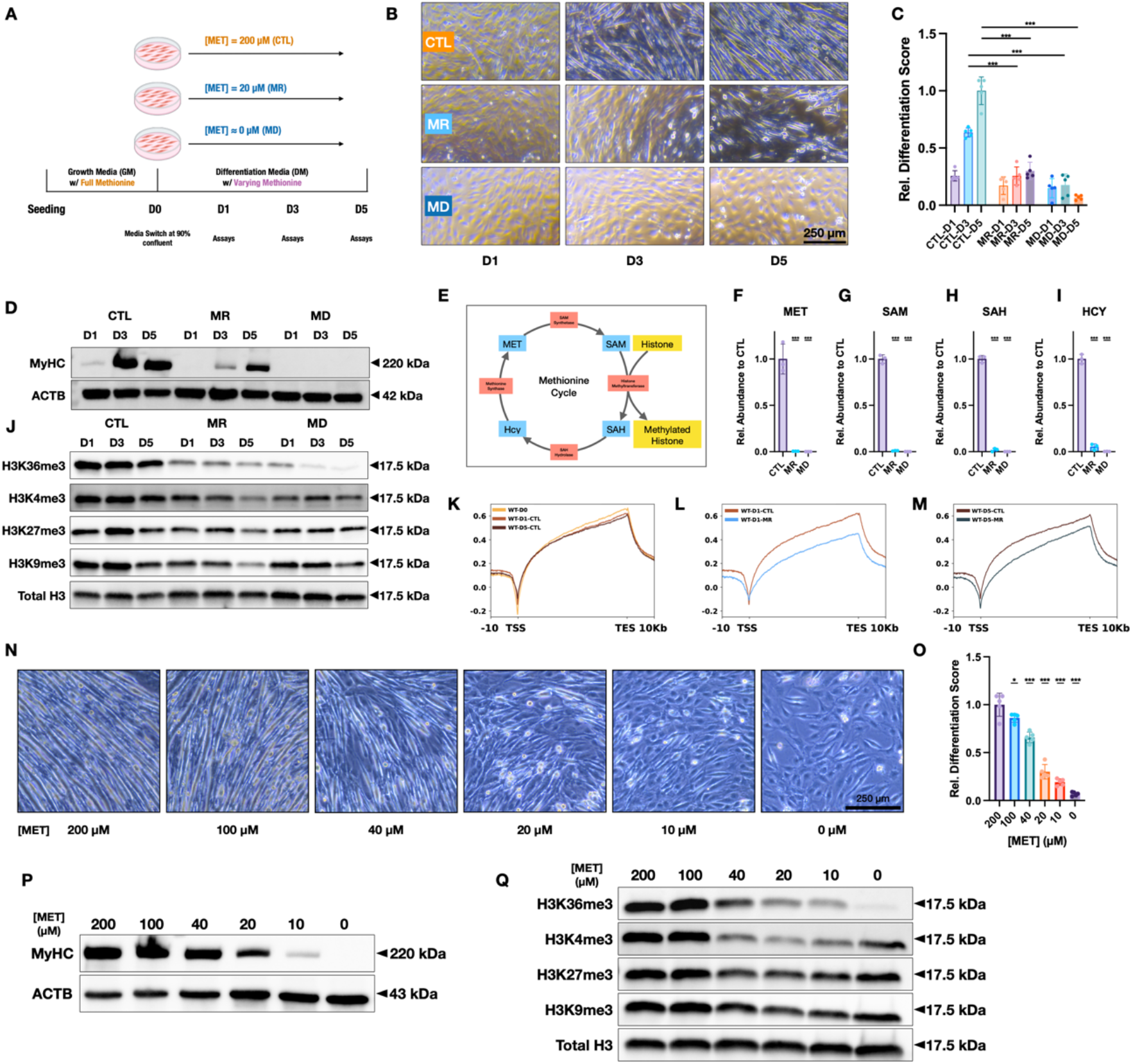
Myogenic differentiation and H3K36me3 are methionine-dependent. (A) Schematic representation of the experimental design: C2C12 cells were transitioned from growth media (GM) with 200 µM methionine (MET) to differentiation media (DM) with methionine either stayed at 200 µM (CTL), restricted to 20 µM (MR) or completely depleted (MD, 0 µM). Imaging and assays were performed on D1, D3 and D5. (B) Representative images of C2C12 cells on Day 1 (D1), Day 3 (D3) and Day 5 (D5) under MET, MR, and MD conditions, displaying the disruption in elongated myotube formation under MR and MD conditions. (C) Quantification of differentiation progress using a custom-trained convolutional neural network to analyze cell images collected by phase contrast microscope. (D) Immunoblot analysis of myosin heavy chain protein (MyHC) expression during differentiation under different methionine conditions. (E) Schematic representation of the methionine cycle with key intermediates. (F-I) Abundance of methionine cycle intermediates under MR and MD relative to CTL. (J) Immunoblot analysis of common histone tri-methylation marks in response to MR and MD. (K) ChIP-Seq profile of H3K36me3 abundance across all genes during normal course of differentiation. TSS: Transcription Start Site, TES: Transcription End Site. (L, M) ChIP-Seq profile of H3K36me3 abundance across all genes in response to MR compared to CTL. (N) Representative images of C2C12 cells on Day 5 (D5) that were kept on differential media with a gradient of methionine concentration. (O) Quantification of differentiation progress on Day 5 (D5) cell images. (P) Immunoblot analysis of myosin heavy chain protein (MyHC) expression on Day 5 (D5) in cells that were kept on differentiation media with a varying methionine concentration. (Q) Immunoblot analysis of common histone tri-methylation marks on Day 5 (D5) in cells that were kept on differentiation media with a varying methionine concentration.

To better understand the interplay between methionine availability, differentiation, and histone methylation, we subjected cells to varying methionine concentrations. The results revealed a dose-dependent relationship between methionine concentration, relative differentiation score, MyHC expression, and H3K36me3 levels (Figures 1N - Q), indicating a methionine dose dependent effect on the phenotypes. Collectively, our data underscore the dose-responsive, methionine-dependent nature of myogenic differentiation and H3K36me3 dynamics.

### Methionine dynamically controls H3K36me3 and myogenic differentiation

Next, we explored the potential reversibility of a methionine restriction-induced decrease in H3K36me3 levels and differentiation impediment. Cells were subjected to either MR or MD conditions over the five-day differentiation window, leading to anticipated disruption in the differentiation process. Following this phase, methionine was reintroduced at the control concentration of 200 µM (Figure 2A). Remarkably, this led to a progressive recovery of the differentiation process in both MR and MD pre-treated cells (Figures 2B, 2C), accompanied by a restoration of H3K36me3 levels (Figure 2D). This result underscores the reversible nature of methionine-dependent H3K36me3 modulation and associated differentiation dynamics. Crucially, the reversible nature of these phenotypes allows us to narrow the scope of the potential underlying mechanisms. Specifically, this reversibility suggests mechanisms that are inherently incompatible with such dynamic behavior, such as increased mutation rates from impaired DNA damage repair systems or widespread chromatin instability stemming from H3K36me3 loss, can be ruled out. Thus, our observations are consistent with transient and reversible shifts in the epigenetic landscape mediated by methionine.

**Figure 2:**
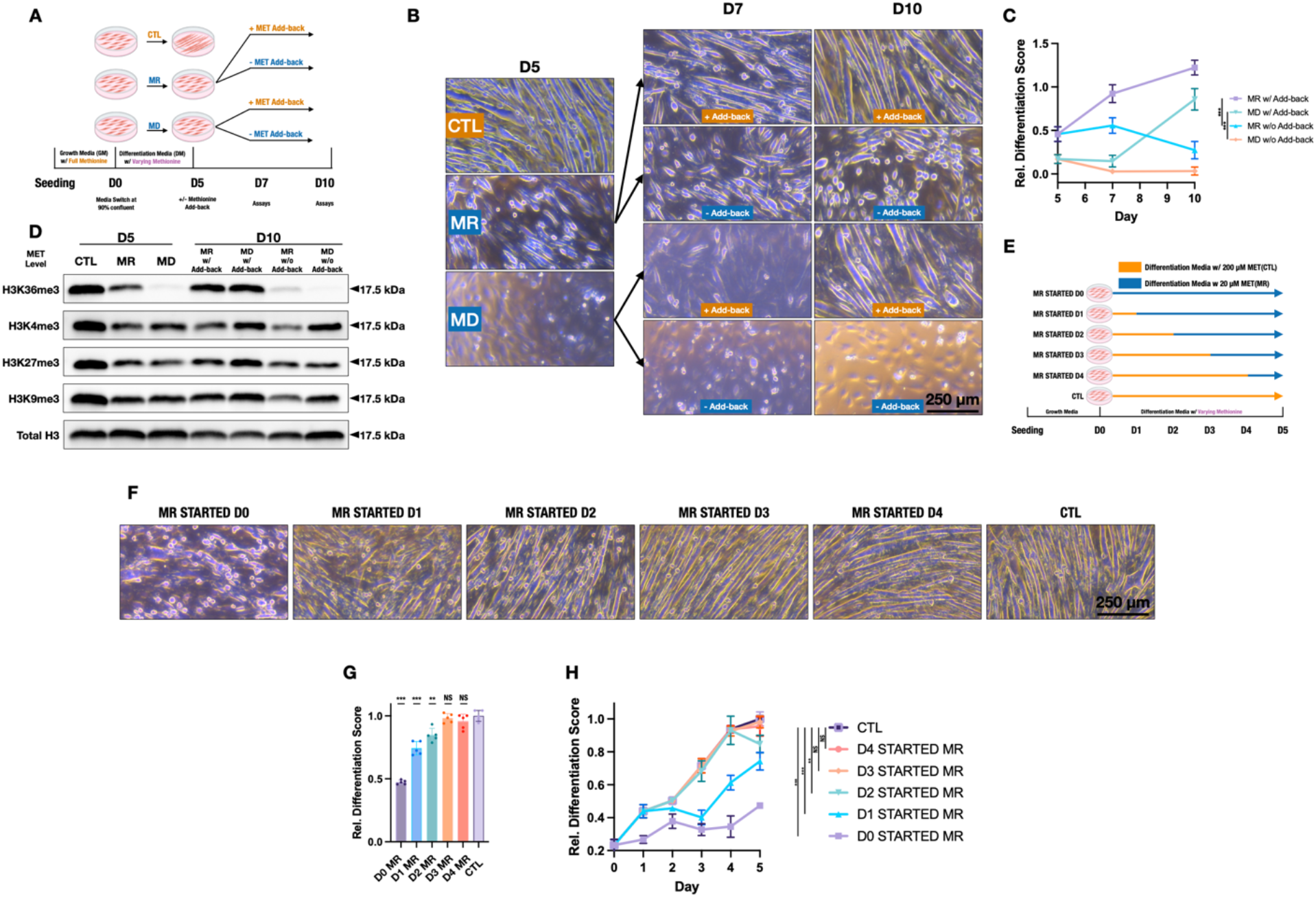
Methionine dynamically controls H3K36me3 and myogenic differentiation. (A) Experimental design depicting the normal 5-day differentiation period with methionine restriction (MR) or depletion (MD), followed by additional 5-day differentiation period with or without reintroduction of methionine at control concentration (200 µM). (B, C) Representative images of C2C12 cells and differentiation score showcasing differentiation recovery in MR and MD pre-treated cells post-methionine add-back. (D) Immunoblot analysis of common histone tri-methylation marks showcasing restoration of H3K36me3 levels upon methionine add-back. (E) Schematic of the methionine cut-off experiment design, where MR was introduced at different time points across the differentiation window. (F, G) Representative images of C2C12 cells and differentiation score at Day (D5) endpoint. (H) Time series plot of differentiation scores demonstrating that initiating MR at D2 has minimal impact on differentiation.

To further elucidate the potential mechanism of MR induced differentiation impediment, we focused on identifying the stage at which methionine is most critical for the differentiation process. To this end, we conducted a methionine cut-off experiment. After allowing cells to reach confluence, MR was introduced at varying time points across the differentiation window to pinpoint when methionine exerts its most pivotal role (Figure 2E). As expected, initiating MR from D0 disrupted the differentiation process (Figure 2F). Yet, intriguingly, introducing MR from D2 onward had minimal effect on differentiation by the end of the five-day period (Figures 2G, 2H). This observation holds important implications. By D2, cells have yet to exhibit the primary morphological changes characteristic of differentiation, namely the formation of elongated myotubes. If a deficiency in methionine as an essential nutrient solely acts by impeding the synthesis of MyHC and other myogenic related proteins - then introducing MR from D2 should mimic the differential impediments observed when MR is applied throughout. Such disparity hints at the possibility that MR chiefly affects differentiation by preventing cells from orchestrating the essential gene expression profiles necessary for myogenic differentiation upon differentiation initiation.

### Myogenic differentiation and H3K36me3 are SAM-dependent

To further dissect the role of methionine in differentiation, we focused on its function as a methyl group donor. This was achieved by targeting the availability of SAM, a key metabolic product of methionine that acts as the primary methyl donor in cells. Given concerns regarding SAM transport in mammalian cells^18^, we opted not for direct SAM supplementation under MR but rather for a strategy to restrict SAM while keeping methionine at control levels. By inhibiting the SAM synthetase, MAT2A, with cycloleucine (cLEU) (Figure 3A)^19^, we noted reductions in the levels of methionine cycle metabolites (Figure 3B, 3D, 3E), except for methionine itself, whose intracellular levels even increased (Figure 3C). Yet, despite sufficient methionine, differentiation disruption akin to MR conditions was observed (Figure 3G, 3H), accompanied by a decrease in H3K36me3 levels (Figure 3F). Taken together, our findings suggest that the ability of methionine to act as a methyl group donor is essential for the observed differentiation outcomes under MR conditions.

**Figure 3:**
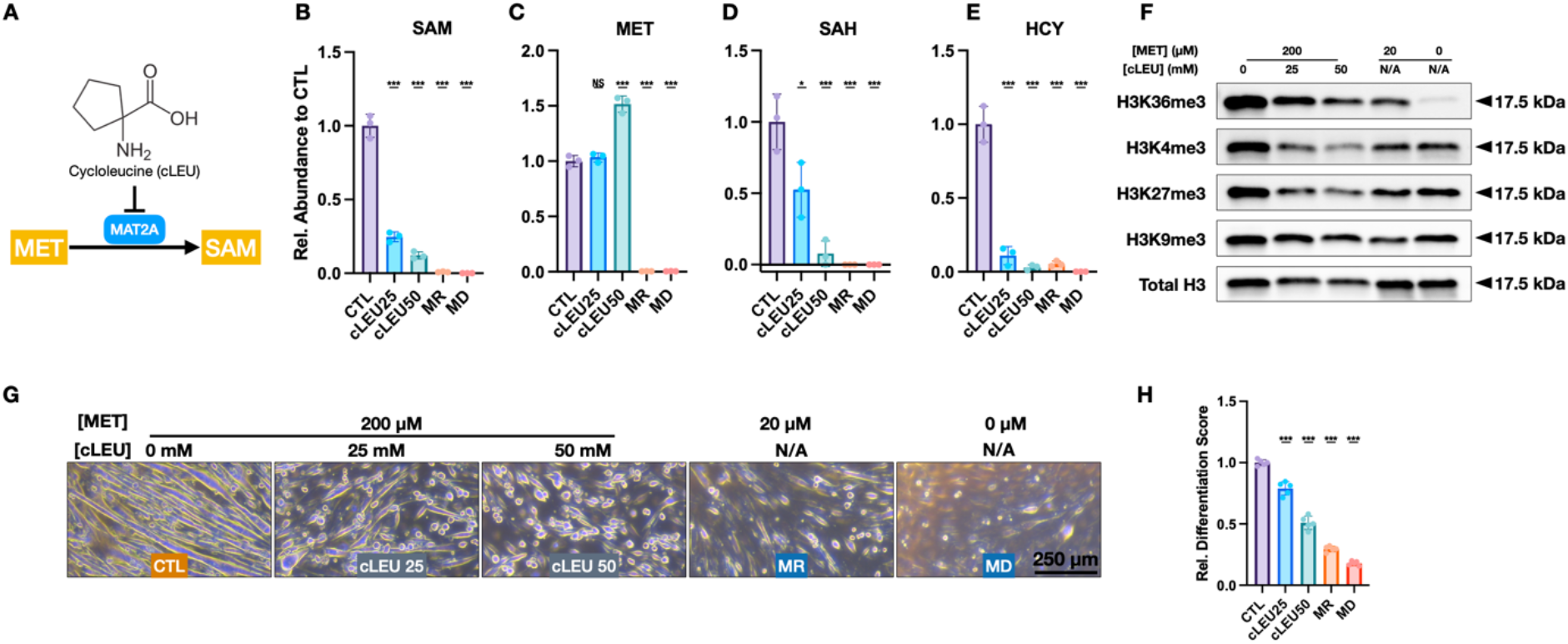
C2C12 Differentiation and H3K36me3 are S-Adenosylmethionine (SAM)-dependent. (A) Schematic representation of SAM synthesis highlighting the SAM synthetase, MAT2A, and its inhibition by cycloleucine (cLEU). (B-E) Abundance of methionine cycle intermediates under cLEU, MR and MD relative to CTL. (F) Immunoblot analysis of common histone tri-methylation marks upon MAT2A inhibition, MR and MD. (G, H) Representative images of C2C12 cells and differentiation score at Day (D5) endpoint.

### Methionine-dependent differentiation acts through SETD2

After establishing the role of SAM in myogenic differentiation, we sought to identify its downstream effectors. The observed decrease in H3K36me3 levels, paired with differentiation defects under MR or SAM synthetase inhibition, directed our attention to the histone methyltransferase known to primarily be responsible for H3K36me3, SETD2^20^. Additionally, we observed that SETD2 levels peak early in and later decrease during differentiation (Figure 4A), consistent with methionine being required at the onset of differentiation (Figure 2F). These observations further suggest that the impact of methionine on differentiation may, in part, operate through SAM and SETD2. To further investigate this, we ectopically-expressed a truncated version of SETD2 (tSETD2)^21^ using a lentivirus system (Figure 4B). The ectopic expression of tSETD2 not only increased H3K36me3 levels under MR conditions (Figure 4C) but also ameliorated the differentiation defects caused by MR evident by cell morphology, differentiation extent and MyHC expression (Figures 4D - F). Conversely, knockout of Setd2, via CRISPR-Cas9 (Figure 4G, Methods), led to a stark reduction in H3K36me3 levels and impaired the differentiation process under standard levels of methionine (Figures 4H - K). Moreover, the Setd2-knockout cells remained indifferent to changes in methionine concentration, even at twice the control or MR levels (Figures 4H - K). Collectively, these findings reinforce the role of SETD2 as a key intermediary in the differentiation process influenced by methionine.

**Figure 4.**
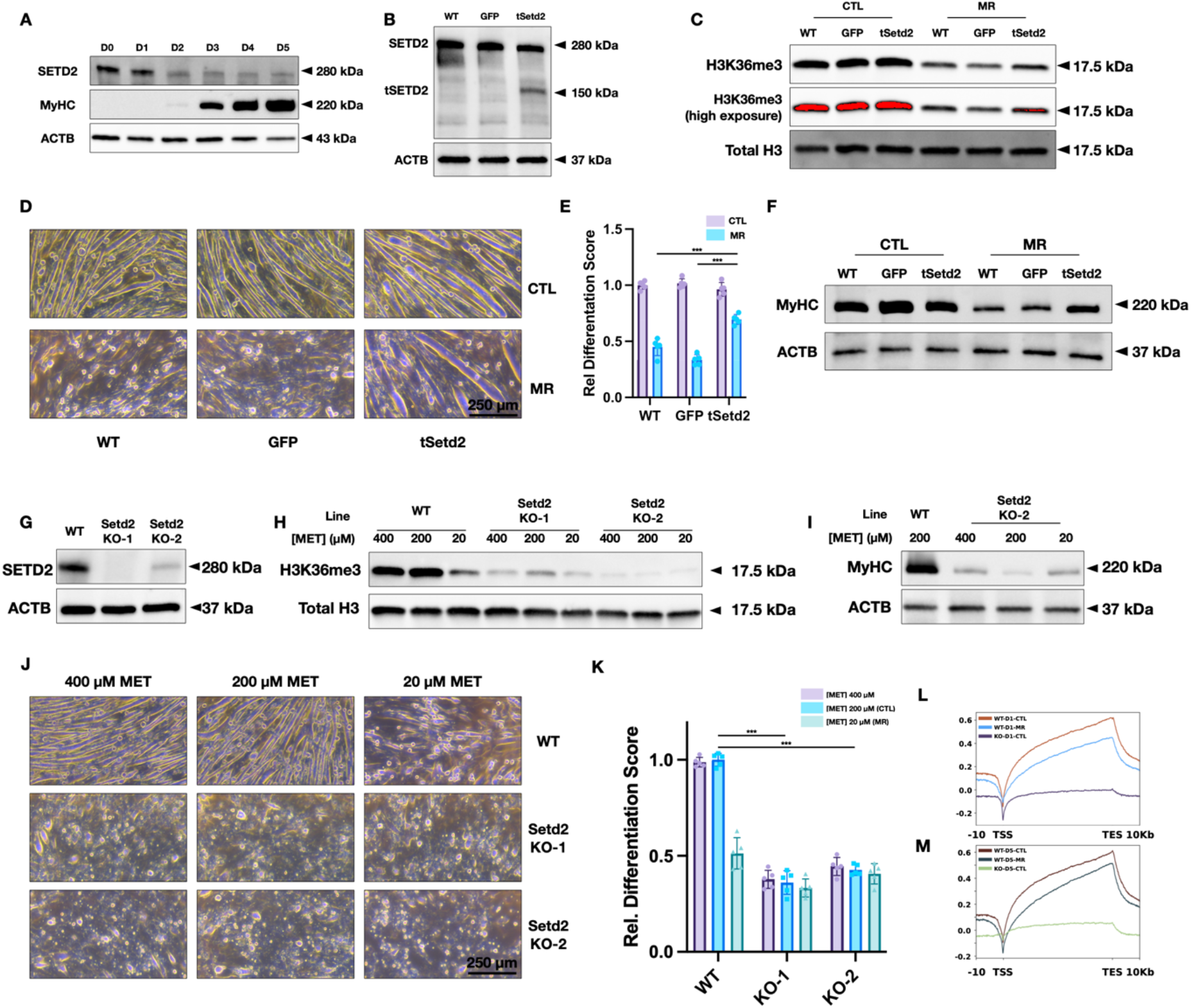
Methionine-dependent differentiation acts through SETD2. (A) Immunoblot analysis of SETD2 levels during the five-day differentiation window. (B) Immunoblot analysis of SETD2 and truncated SETD2 (tSETD2) levels in WT, WT-GFP and WT-tSetd2 cells. (C) Immunoblot analysis of H36K36me3 in WT, WT-GFP and WT-tSetd2 cells under CTL and MR conditions. (D, E) Representative images of C2C12 cells and differentiation score at Day (D5) endpoint, showing improvements in differentiation progress in WT-tSetd2 cells. (F) Immunoblot analysis of myosin heavy chain protein (MyHC) expression at Day (D5) endpoint. (G) Immunoblot analysis of SETD2 levels in WT, Setd2 knockout cell lines. (H) Immunoblot analysis of H36K36me3 in WT and Setd2 knockout cell lines under 2X, CTL and MR conditions. (I) Immunoblot analysis of myosin heavy chain protein (MyHC) expression at Day (D5) endpoint in WT and Setd2 knockout cell lines under 2X, CTL and MR conditions. (J, K) Representative images of C2C12 cells and differentiation score at Day (D5) endpoint, showing disruption in differentiation in Setd2 knockout cell lines. (L, M) ChIP-Seq profile of H3K36me3 abundance across all genes in response to MR and Setd2 knockout compared to CTL.

### Methionine restriction preferentially reduces H3K36me3 levels at differentiation related genes

Recognizing that the influence of methionine on differentiation can operate through SETD2, we sought to understand the role of its product, H3K36me3, in methionine-dependent differentiation. For this, we analyzed the H3K36me3 ChIP-Seq data of myogenic-related genes that are the targets of key myogenic transcription factors (Table 1). Contrary to the relatively stable aggregated H3K36me3 levels seen across all genes over the 5-day differentiation window seen earlier (Figure 1K), levels of H36K3me3 at myogenic-related genes displayed steady increase, peaking on D5 (Figure 5A). Predictably, MR also diminished the H3K36me3 levels at these genes (Figures 5B, 5C). Notably, effects of MR on H3K36me3 levels varied among different genes. For instance, MR stunted the incremental increase of H3K36me3 levels at Myog and Mef2c, and negated the consistent levels of H3K36me3 seen at Myod1. In contrast, Myh genes displayed subtler MR effects. Some genes that are not directly tied to myogenic differentiation, like Setd2, exhibit no significant changes in H3K36me3 levels in response to MR (Figure 5D - H, suggesting a preferential effect of MR on differentiation-related genes.

**Figure 5.**
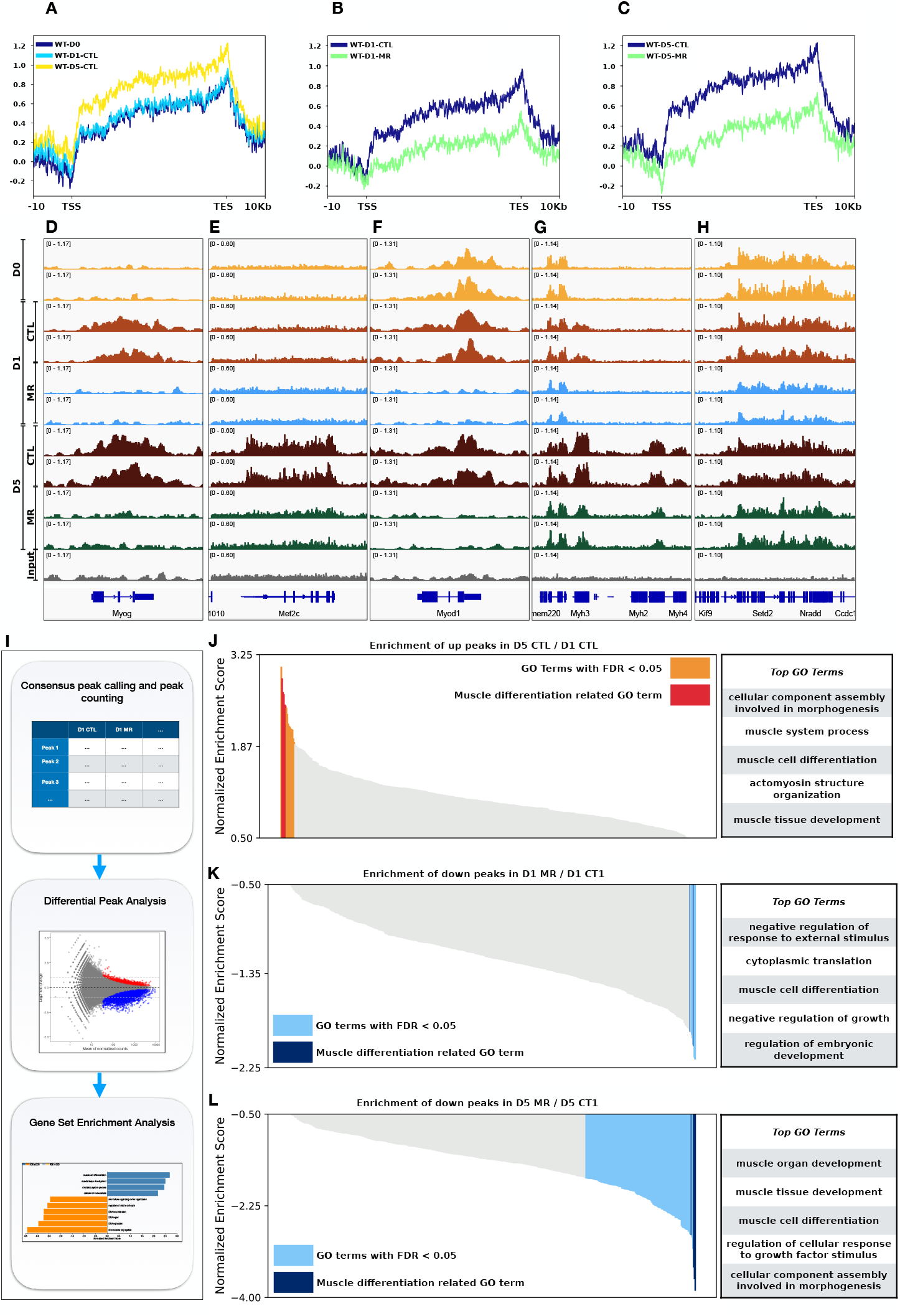
Methionine Restriction Preferentially Diminishes H3K36me3 Levels at Differentiation-Related Genes. (A) ChIP-Seq profile of H3K36me3 abundance across myogenic differentiation-related genes during normal course of differentiation. (B, C) ChIP-Seq profile of H3K36me3 abundance across myogenic differentiation-related genes in response to MR compared to CTL. (D-H) Gene-specific H3K36me3 profiles under CTL and MR conditions. (D) Myog (E) Mef2c (F) Myod1 (G) Myh genes (H) Setd2 (I) Schematic representation of the workflow for differential ChIP peak analysis of H3K36me3 under different methionine conditions. (J) Gene Set Enrichment Analysis (GSEA) of genes displaying increased H3K36me3 levels in a D5 vs. D1 comparison at control methionine levels, with muscle differentiation related Gene Ontology (GO) terms dominate the top of the list. (K, L) GSEA of genes with diminished H3K36me3 peaks under MR: (K) D1 comparison (L) D5 comparison, with muscle differentiation-related GO terms dominate the top of the lists.

To further investigate this, we analyzed the differential ChIP peaks of H3K36me3 under varied conditions, followed by gene set enrichment analysis (Figure 5I). A D5 vs D1 comparison at standard methionine levels revealed GO terms related to muscle differentiation being enriched in genes that saw increased H3K36me3 levels (Figure 5J). Conversely, peaks diminished by MR at either D1 or D5 also showed an enrichment in muscle differentiation-related GO terms with D5 exhibiting more myogenic differentiation-related GO than D1 (Figures 5K, 5L). Collectively, these findings indicate that MR selectively reduces H3K36me3 levels at genes vital for myogenic differentiation.

### Methionine restriction and Setd2 knockout preferentially reduce chromatin accessibility at differentiation related genes

To further explore the implications of methionine restriction on the broader epigenetic landscape, we performed Assay for Transposase-Accessible Chromatin using sequencing (ATAC-Seq) on both WT and KO cells under CTL and MR condition. We proceeded with a differential peak analysis followed by Genomic Regions Enrichment of Annotations (GREAT) analysis^22^ (Figure 6A). Intriguingly, in WT cells, ATAC-Seq peaks that showed increased intensity from D0 to D1 were found to be enriched with GO terms pertinent to muscle differentiation (Figure 6B). In the D1 MR versus D1 CTL comparison, peaks with diminished intensity showed a similar enrichment in muscle differentiation-related GO terms (Figure 6C). Similarly, a D1 KO CTL to D1 WT CTL comparison unveiled peaks with reduced intensity again enriched in muscle differentiation-related GO terms (Figure 6D). Further, on examining ATAC-Seq peaks that were reduced in response to MR, we noticed these peaks were also lowered in the KO cells under CTL condition. Crucially, subjecting KO cells to MR did not further diminish their levels (Figure 6E). Collectively, our findings suggest that the impact of MR on epigenetic landscape, as judged by chromatin accessibility, is mediated, at least in part, through Setd2.

**Figure 6:**
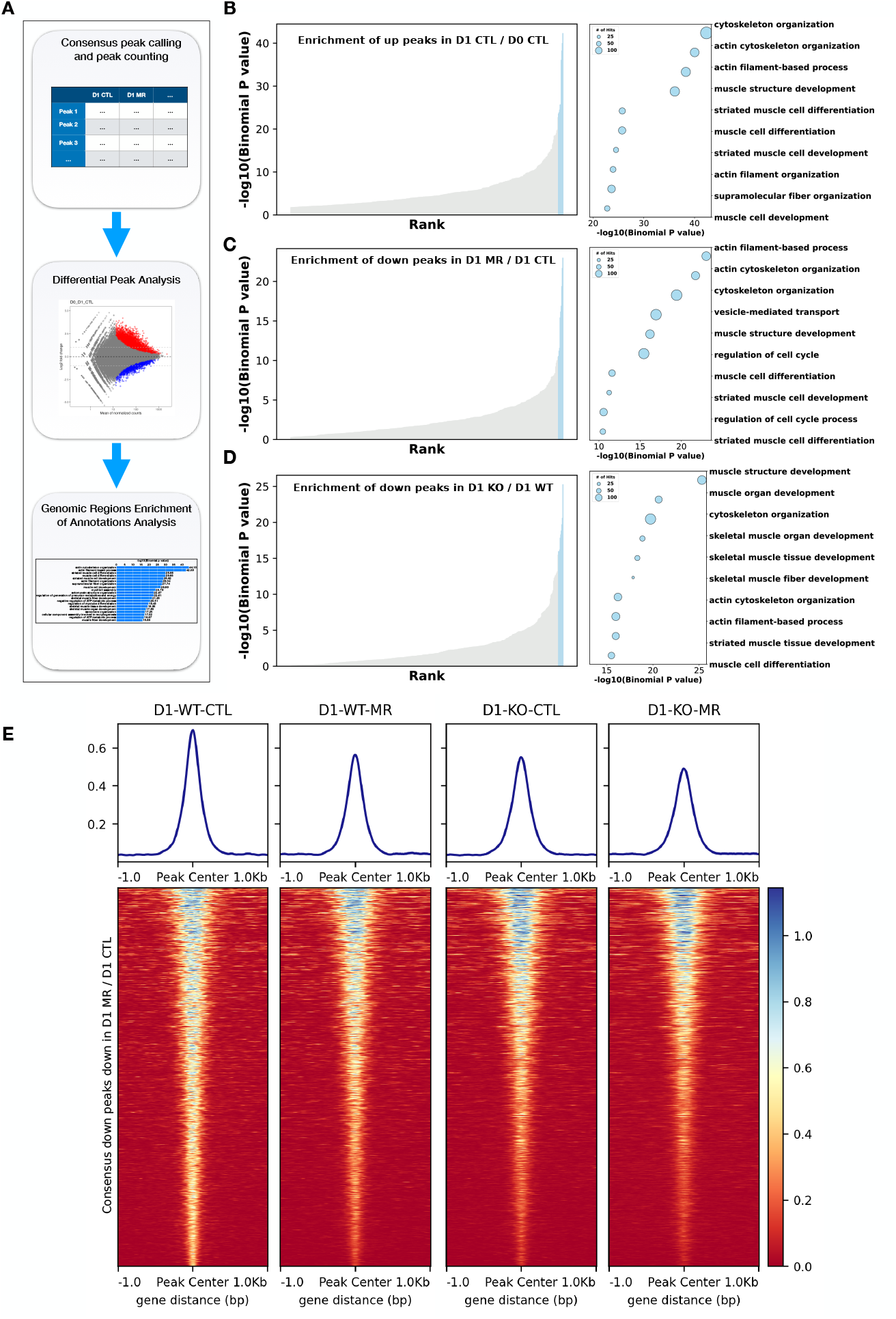
Methionine restriction and Setd2 knockout preferentially reduce chromatin accessibility at differential related genes. (A) Schematic representation of the workflow for differential ATAC-Seq peak analysis under different methionine conditions on WT and Setd2 knockout lines. (B) Genomic Regions Enrichment of Annotations (GREAT) analysis of ATAC-seq peaks that increased in intensity from Day 0 (D0) to Day 1 (D1) in WT cells. Muscle differentiation-related GO terms are predominantly enriched. (C) GREAT analysis of ATAC-seq peaks with diminished intensity comparing Day 1 under MR conditions to Day 1 under CTL conditions in WT cells. Muscle differentiation-related GO terms are predominantly enriched. (D) GREAT analysis of ATAC-seq peaks with diminished intensity between KO cells under CTL conditions and WT cells under CTL conditions on Day 1. Muscle differentiation-related GO terms are predominantly enriched. (E) ATAC-seq profiles and heatmaps of peaks that are diminished in response to MR compared to CTL in WT, and Setd2 knockout lines under CTL and MR conditions. Notably, applying MR to KO cells did not lead to a further reduction in profile peak intensities.

## Discussion

The interplay between metabolism and epigenetics has increasingly come under the spotlight in recent years. While this connection is established, the critical details of how specific metabolic processes and metabolites fine-tune the epigenetic landscape are yet to be thoroughly elucidated. Historically, the study of epigenetic processes primarily rested upon genetic manipulations of chromatin or nucleic acid factors. Yet, the enzymatic kinetics of certain modifiers, such as methyltransferases, present opportunities for regulation via substrate concentration changes, enabled through dietary measures and such as methionine restriction and other means of altering metabolism.

Our study revealed the dependency of H3K36me3 and myogenic differentiation on methionine, offering insights into the dynamics and reversibility of these dependencies. Further, we explored the mechanism behind this methionine dependency by establishing the role of SAM and Setd2 as instrumental links in the causal chain connecting methionine to differentiation, suggesting that SETD2 might act as a sensor for methionine in certain settings. Further establishing the link to gene regulation, our H3K36me3 ChIP-Seq and ATAC-Seq data provide additional insights on how methionine and Setd2 ultimately contribute to myogenic differentiation.

Establishing a direct causal chain spanning methionine to SAM, Setd2, H3K36me3, and finally differentiation, is complex owing to various confounding elements. Every component in this pathway has multifaceted roles. For instance, methionine serves functions beyond methyl group donation, playing a part in protein synthesis and redox balance^23^. SAM is used by a plethora of methyltransferases, while Setd2 has diverse functions, including interactions with RNA Pol II^24^. Fully deciphering the contribution of these intertwined elements is difficult. Nevertheless, our efforts established the methionine-SAM-Setd2 axis’ significance in cell differentiation. Furthermore, our results demonstrated that cellular and genomic outcomes that traditionally achieved via genetic manipulations, such as Setd2 knockout, can also be partially if not largely recapitulated by altering the nutrient environment of the cells, opening new directions of investigations. Overall, this study establishes the direct sensing of methionine to chromatin in the context of myogenic differentiation with specific components, Setd2 and H3K36me3, both of which are known to be involved in key process in development and cancer, contributing to the regulation of epigenetic landscape.

## Methods

### Cell culture

C2C12 cells were procured from the Duke Cell Culture Facility. They were maintained at 37°C and 5% CO2 on growth media (GM) for C2C12 comprised of DMEM (Gibco 11965092), 10% heat-inactivated fetal bovine serum (Sigma-Aldrich F2442), and 1X penicillin-streptomycin (Gibco 15140163). For differentiation induction, cells were seeded at a density of 1.25E5 cells/well in a 6-well plate and allowed to proliferate for 48 hours in GM to reach confluence. Post confluence, the culture medium was switched to differentiation media (DM), which consisted of DMEM without methionine, cystine-2HCl, and glutamine (Gibco 21013024), supplemented with 2% heat-inactivated horse serum (Hyclone SH30074.03), 1X pen/strep (Gibco 15140163), and 1 µM insulin (Sigma-Aldrich I6634). Both cystine-2HCl and glutamine were reintroduced to levels found in standard DMEM. Depending on the experimental conditions, methionine was either added to a concentration of 200 µM (for control) or to other specified concentrations. The differentiation process was allowed to proceed for 5 days, with media changes occurring every 48 hours.

### Microscopy

Images of C2C12 cells were acquired using a Leica DM IL LED microscope, which was outfitted with a Leica MC170HD camera. Imaging was performed using 20x, 10x, and 4x objectives, and images were captured and processed using the LAS EZ software.

### Differentiation Quantification

A convolutional neural network (CNN) was trained using the ResNet-18 architecture with PyTorch^25,26^. We modified ResNet-18 by replacing its final fully connected layer to yield a single continuous output value. 64 images, captured using the 10x objective, were obtained each day from Day 0 (D0) to Day 5 (D5) for cells exposed to differentiation media (DM) containing full methionine. To augment our dataset, each of these images was subdivided into five sub-images. For labeling, each image was assigned a numeric value corresponding to the day; e.g., an image of a Day 3 (D3) cell received a label of 3.0. Out of the 320 augmented images obtained each day, 270 of them were used for training while the remaining 50 of them served as the test set. The CNN was trained for 15 epochs using a learning rate of 1e-4, utilizing the Mean Squared Error Loss (nn.MSELoss) as the criterion, and Adam as the optimizer. To ascertain the differentiation score of a given image relative to image of cells on a control condition—in which cells differentiated over an identical duration—each image was partitioned into five sub-images and assessed five times. Subsequent results were normalized to the average score of the control group.

### Histone extraction

Cells were washed twice with room-temperature PBS before being scraped into ice-cold Triton extraction buffer (TEB) containing 0.5% Triton X-100, 0.2% NaN3, and 2 mM PMSF in PBS. The suspension was transferred to a 1.5 ml centrifuge tube and briefly vortexed, followed by a 10-minute incubation on ice. The cells were then centrifuged at 12,000 g at 4°C for 10 minutes. After discarding the supernatant, the cell pellet was resuspended in 500 µl of TEB and centrifuged again using the same parameters. The supernatant was removed, and the cell pellet was resuspended in 500 µl of 0.2N HCl, which was then incubated at 4°C overnight. The next day, the samples were centrifuged at 12,000 g at 4°C for 10 minutes. The supernatant was collected, neutralized with 2M NaOH, and their concentrations were determined using a BCA assay.

### Total protein extraction

Cells were rinsed twice with room-temperature PBS and subsequently scraped into ice-cold RIPA buffer (Sigma-Aldrich R0278) supplemented with a 1x Protease Inhibitor Cocktail (Roche 11836170001). After transferring the cell suspension to a 1.5 ml centrifuge tube, it was briefly vortexed and then incubated on a nutator at 4°C for 25 minutes. Following incubation, the cells were centrifuged at 12,000 g at 4°C for 25 minutes. The supernatant was carefully collected, and protein concentrations were quantified using a BCA assay.

### Immunoblot

Samples were diluted to the desired concentrations (4 mg/well for histone and 15 mg/well for total protein extracts) using water and combined with Laemmli sample buffer containing 10% 2-Mercaptoethanol. This mixture was heated at 95°C for 5 minutes. The samples were subsequently subjected to electrophoresis on 12% SDS-PAGE gels for histone samples and 4-15% SDS-PAGE gels for total protein extracts, after which they were transferred onto PVDF membranes (Bio-Rad 1704156).The membranes were blocked using 5% BSA (Rockland BSA-50) in TBS-T buffer for an hour, then probed overnight at 4°C with primary antibodies, all diluted 1:1000: Myosin Heavy Chain (R&D System mAB4470), ActB (CST 8457T), Histone H3 (CST 96C10), H3K36me3 (Active Motif 61101), H3K27me3 (Active Motif 61018), H3K9me3 (Abcam ab8898), H3K4me3 (Active Motif 39060), and Setd2 (CST E4WQ8). After overnight incubation, membranes were washed three times with TBS-T, each for 5 minutes. This was followed by a one-hour incubation at room temperature with peroxidase-conjugated Rabbit or Mouse secondary antibodies (Rockland 610-1302 and 611-7302) at a dilution of 1:10000. Post-incubation, membranes underwent another set of three washes with TBS-T, each lasting 5 minutes. Membranes were then exposed to the ECL substrate kit (Pierce 34850 and 34095) for 3 minutes before imaging with the ChemiDoc Touch Imaging System (Bio-Rad).

### Metabolites extraction

Cell culture medium was quickly removed before adding 1 ml of an 80% methanol/water solvent (Fisher Scientific A456 and W6), cooled to −80°C, to each well of the six-well plates. These plates were then chilled at −80°C for 15 minutes. After this, the cells were scraped into the solvent while on dry ice, and the extracts were centrifuged at 20,000g at 4°C for 10 minutes. The solvent was then evaporated using a speed vacuum and the dried metabolites pallets were stored at -80°C.

### Liquid chromatography

Dried metabolites pallets were reconstituted in 15 μl of H2O, followed by the addition of 15 μl of a 1:1 v/v methanol/acetonitrile mix (Fisher Scientific A456 and A955). The samples were then centrifuged at 20,000g at 4°C for 10 minutes, after which the supernatants were moved to LC vials, preparing them for a 3 μl HPLC injection. The compounds were separated at room temperature using an XBridge amide column (Waters 100 × 2.1 mm i.d., 3.5 μm;) on a Thermo Scientific UltiMate 3000 UHPLC system. Mobile phases and flow rate was configured as described in previous study^27^.

### Mass spectrometry

The Q Exactive Plus mass spectrometer (Thermo Fisher Scientific) was equipped with a heated electrospray ionization probe and configured with the following settings: heater temperature at 120°C; sheath gas flow rate at 30; auxiliary gas at 10; and sweep gas at 3. The spray voltage was set to 3.6 kV in positive mode and 2.5 kV in negative mode. The capillary temperature and S-lens were maintained at 320°C and 55, respectively. Scans were conducted across a range of 60 to 900 m/z at a resolution of 70,000. The automated gain control (AGC) was targeted at 3E6 ions.

### Peak extraction and metabolomics data analysis

The acquired raw LC-MS data were analyzed using Thermo Scientific Sieve 2.0 software, with peak alignment and detection executed as per the manufacturer’s guidelines. For targeted metabolite analysis in positive mode, a frame seed encompassing 194 metabolites was utilized, whereas a frame seed comprising 262 metabolites was adopted for the negative mode. The m/z width was set at 10 ppm for both modes. Subsequently, peak integration values were exported. Sample values were normalized based on the mean value of the control group.

### Quantitative PCR

RNA was extracted using the Qiagen RNeasy Micro Kit in accordance with the manufacturer’s instructions. The quality and quantity of the RNA were determined using NanoDrop (Thermo Scientific). Subsequently, 1 µg of RNA was reverse transcribed into single-stranded complementary DNA utilizing the SuperScript III First-Strand Synthesis SuperMix for qRT-PCR from Invitrogen. Quantitative analysis was carried out on a CFX384 system (Bio-Rad) using the iQ SYBR Green Supermix (Bio-Rad) as directed by the manufacturer.

### Lentiviral transduction

Lentivirus containing GFP or truncated Setd2, obtained from Vectorbuilder, was used to transduce C2C12 cells. Cells were seeded in a 12-well plate at a density of 0.45E5 cells per well and allowed to proliferate for 24 hours, reaching approximately 1E5 cells per well. For the transduction, 100 µl of lentivirus (1E8 TU) was combined with 300 µl of fresh growth media containing polybrene at 5 µg/ml. After removing the existing cell culture medium, cells were cultured with the 400 µl of lentivirus-containing media for 24 hours. The following day, cells were transferred to a 10 cm culture dish. Subsequently, puromycin was added at a concentration of 3 µg/ml to initiate the selection process. After 24 hours, surviving cells were relocated to a 12-well plate without puromycin. These cells were then incrementally transferred first to a 6-well plate and later to a 10 cm plate. A final round of selection was done by reintroducing puromycin. The expression levels of tSetd2 and GFP were then confirmed using a fluorescent microscope and immunoblotting.

### CRISPR Knockout of Setd2

CRISPR Cas-9 mediated *Setd2* knockout in C2C12 cells was performed by the Duke Functional Genomics Core. Three sgRNAs targeting *Setd2* were designed using ChopChop and ordered as modified synthetic sgRNAs from Synthego^28^. 5 × 10e4 C2C12 cells were electroporated with 7.5 pmol TrueCut Cas9 protein v2 (ThermoFisher Scientific) complexed with 22.5 pmol sgRNA using the Neon system (ThermoFisher Scientific) with the following settings: 1650 V, 10 ms, 3 pulses. Cells were recovered and expanded, followed by PCR sequencing and immunoblotting analysis to determine KO efficiencies. Based on this analysis, knockout clones were subsequently isolated from the pools generated with sgRNA #1 (GTCGGTCCGAAAGAGATCGA) and sgRNA #3 (GCATTCGCTTAATATCCCGG) by plating at limited dilution in 96 well plates. Clones were expanded for approximately 10-14 days before screening by PCR-sequencing to confirm out-of-frame insertions or deletions.

### ChIP-Seq and data analysis

Samples were sent to Active Motif (Carlsbad, CA) for ChIP-Seq. Active Motif prepared chromatin, performed ChIP reactions, generated libraries, and sequenced the libraries. In brief, C2C12 cells were fixed in formaldehyde. Chromatin was isolated by adding lysis buffer, followed by disruption with a Dounce homogenizer. Lysates were sonicated and the DNA sheared with Active Motif’s EpiShear probe sonicator (cat# 53051). Genomic DNA (Input) was prepared by treating aliquots of chromatin with RNase, proteinase K and heat for de-crosslinking, followed by SPRI beads clean up (Beckman Coulter) and quantitation by Clariostar (BMG Labtech). Extrapolation to the original chromatin volume allowed determination of the total chromatin yield.

An aliquot of chromatin, 30 ug chromatin (750 ng of Drosophila chromatin for spike-in for H3K36me3^29^) was precleared with protein G agarose beads (Invitrogen). Genomic DNA regions of interest were isolated using 4ul of H3K36me3 (Active Motif cat# 61101, lot# 22264149) and 0.4ug of antibody against H2Av (Active Motif cat# 61686). Complexes were washed, eluted from the beads with SDS buffer, and subjected to RNase and proteinase K treatment. Crosslinks were reversed by incubation overnight at 65°C, and ChIP DNA was purified by phenol-chloroform extraction and ethanol precipitation.

Quantitative PCR (qPCR) reactions were carried out in triplicate on specific genomic regions using SYBR Green Supermix (Bio-Rad). The resulting signals were normalized for primer efficiency by carrying out qPCR for each primer pair using Input DNA.

Illumina sequencing libraries (a custom type, using the same paired read adapter oligonucleotides^30^) were prepared from the ChIP and Input DNAs on an automated system (Apollo 342, Wafergen Biosystems/Takara). After a final PCR amplification step, the resulting DNA libraries were quantified and sequenced on Illumina’s NovaSeq 6000 (75 nt reads, single end).

The resulting sequence data were processed using the nf-core/chipseq pipeline^31^. Targets of key myogenic transcription factors were queried from TRRUST v2^32^. deepTools2^33^ were used for generating profile plots, while the Integrative Genomics Viewer was used for track visualization^34^. For the analysis of differential peaks, DESeq2^35^ was used. Gene Set Enrichment Analysis was subsequently performed using WebGestalt^36^.

### ATAC-Seq and data analysis

Cryopreserved C2C12 cells were sent to Active Motif to perform the ATAC-Seq assay. The cells were then thawed in a 37°C water bath, pelleted, washed with cold PBS, and tagmented as previously described^37^, with additional modifications^38^. Briefly, cell pellets were resuspended in lysis buffer, pelleted, and tagmented using the enzyme and buffer provided in the ATAC-Seq Kit (Active Motif). Tagmented DNA was then purified using the MinElute PCR purification kit (Qiagen), amplified with 10 cycles of PCR, and purified using Agencourt AMPure SPRI beads (Beckman Coulter). Resulting material was quantified using Qubit (Invitrogen) and TapeStation (Agilent), then sequenced with PE42 sequencing on the NovaSeq 6000 sequencer (Illumina).

The resulting sequencing data were processed using the nf-core/atacseq pipeline^31^. deepTools2^33^ were used for generating profile plots and heatmaps. For the analysis of differential peaks, DESeq2^35^ was used. Genomic Regions Enrichment of Annotations (GREAT) analysis was subsequently performed using GREAT online server^22,39^.

### Statistical analysis

Independent sample T-tests were conducted using the ttest_ind function from the SciPy stats package^40^. Significance levels are denoted as follows: * for p < 0.05, ** for p < 0.01 and *** for p<0.001. Bar graphs depict the mean, with the error bars representing the standard deviation.

### Generative AI

During the preparation of this manuscript, the authors employed Bard and ChatGPT for language refinement to improve clarity and readability, as well as for assistance with specific aspects of code implementation and debugging. After using these tools, the authors thoroughly reviewed and revised the content as necessary. We take full responsibility for the final content of this publication.

## Data availability

The dataset for ChIP-seq and ATAC-seq experiments generated in this study is publicly available at the NCBI Gene Expression Omnibus (accession no: GSE248198 and accession no: GSE248197). The code for the differentiation quantification program is publicly available on GitHub. (https://github.com/LocasaleLab/DiffQuant)

## Acknowledgement

The authors extend their gratitude to all members of the Locasale Lab, with special thanks to Dr. Juan Liu, Dr. Shiyu Liu, Dr. Annamarie E. Allen and Luis G. Vergara for their insightful discussions and technical assistance. We are grateful to the National Institutes of Health (R01CA193256 to JWL) and the American Cancer Society (129832-RSG-16-214-01-TBE to JWL) for their generous funding. Additionally, we acknowledge the support for CRISPR-Cas9 provided by the Duke Functional Genomics Core.

